# Reduced plasma hexosylceramides in frontotemporal dementia are a biomarker of white matter integrity

**DOI:** 10.1101/2025.02.17.638741

**Authors:** Oana C. Marian, Sophie Matis, Carol Dobson-Stone, Woojin S. Kim, John B. Kwok, Olivier Piguet, Glenda M. Halliday, Ramon Landin-Romero, Anthony S. Don

## Abstract

**INTRODUCTION:** Blood biomarkers are needed to facilitate new therapeutic trials and improve management of behavioural variant frontotemporal dementia (bvFTD). Since altered white matter integrity is characteristic of bvFTD, this study aimed to determine if plasma levels of myelin-enriched glycolipids are altered in bvFTD and correlate with white matter integrity.

**METHODS:** Nineteen glycolipids were quantified in bvFTD (n=31) and control (n=26) plasma samples. White matter integrity was assessed using MRI-derived fibre tract density and cross-section (FDC).

**RESULTS:** Eleven lipids were significantly lower in bvFTD compared to control subjects, seven were inversely correlated with disease duration, and twelve were positively correlated with cognitive performance, with C22:0 hexosylceramide most strongly correlated. FDC was lower in frontotemporal white matter tracts of bvFTD compared to control subjects, and plasma C22:0 hexosylceramide was significantly correlated with FDC in these tracts.

**DISCUSSION:** Circulating glycolipids may be a valuable biomarker of myelin integrity and disease progression in FTD.

## 1 Introduction

Frontotemporal dementia (FTD) is the second-most common cause of younger onset dementia, associated with progressive atrophy of the frontal and temporal lobes.^1^ Behavioural variant FTD (bvFTD) is the most common form, characterised by cognitive, personality and behaviour changes.^1,2^ A family history is reported for nearly half of all bvFTD cases, with the most common gene variants being hexanucleotide repeats in the *C9ORF72* gene and heterozygous mutations in the *GRN* or *MAPT* genes.^3^ Diagnosis is based on a combination of clinical examination, neuroimaging and genetic screening.^1,2^ Sensitive and reliable biomarkers are needed for improved diagnosis of bvFTD and to facilitate the development of effective therapeutics.

Neuroimaging using traditional diffusion tensor imaging metrics has shown early and progressive changes to white matter integrity affecting the uncinate fasciculus, cingulum, and corpus callosum in bvFTD.^4-7^ Recent advances in diffusion-weighted imaging have enabled more biologically interpretable tract-based measurements of white matter integrity,^8^ with the potential to reveal associations between molecular and microstructural changes.

Myelin is a particularly lipid-rich structure and approximately 20% of myelin lipid is galactosylceramide (GalCer), which is relatively unique to myelin.^9^ Its structural isomer glucosylceramide (GluCer) is more abundant in peripheral organs, where GalCer is generally thought to be either absent or present only at trace levels.^10^ Since GluCer and GalCer are mass isomers and cannot be distinguished with conventional reverse-phase liquid chromatography-tandem mass spectrometry (LC-MS/MS) lipidomic analyses, they are collectively referred to as hexosylceramide (HexCer). Lipidomic analysis has demonstrated pronounced loss of myelin-enriched sphingolipids including HexCer in frontal white matter of familial bvFTD cases,^11^ however sporadic bvFTD cases were not examined. We recently developed a hydrophilic interaction chromatography (HILIC) LC-MS/MS method to enable GalCer and GluCer separation and quantification.^12^ This study aimed to determine firstly if levels of GalCer or GluCer species are altered in the plasma of people with bvFTD; and secondly, if these lipids are correlated with fibre-specific measures of white matter integrity.

## 2 Methods

### 2.1 Participants

Study participants (Table 1) were recruited through FRONTIER, the frontotemporal dementia research clinic in Sydney. Participants are reviewed annually, following a comprehensive clinical assessment, cognitive examination, brain MRI and informant report. Diagnosis of bvFTD was established at baseline according to current clinical diagnostic criteria^2^, and confirmed at the latest available follow-up. The Frontotemporal dementia Rating Scale (FRS) was used to measure disease severity. Overall cognition was assessed using either the Addenbrooke’s Cognitive Examination-revised (ACE-R) or ACE-III. Before analyses, ACE-III scores were converted to ACE-R where necessary.^13^ Healthy controls were recruited from the FRONTIER database or from the community. Controls scored ≥ 88/100 on ACE-R or ACE-III and 0 on the Clinical Dementia Rating scale.^14^ Exclusion criteria for all participants included lifelong history of psychiatric disease, presence of other neurodegenerative conditions or neurological disorders, and history of substance abuse. Genetic abnormalities in the *C9ORF72, GRN*, and *MAPT* genes were identified using whole exome sequencing.^15^

**Table 1.** Cohort demographic and clinical data.

| Characteristic | bvFTD (*n*=31) | Control (*n*=26) | Statistic | *P* |
| --- | --- | --- | --- | --- |
| Sex (Female, Male) | 13, 18 | 14, 12 | 0.81* | .37 |
| Age at blood draw (years) ‡ | 65.10 ± 6.81 | 69.67 ± 6.16 | 2.63<sup>†</sup> | **0.01** |
| Education (years) | 12.16 ± 3.08 | 14.24 ± 2.63 | 2.71<sup>†</sup> | **0.009** |
| ACE-R<sup>§</sup> | 71.13 ± 20.07 | 95.37 ± 2.67 | 4.58<sup>†</sup> | **< 0.0001** |
| Disease duration (years) ¶ | 5.82 ± 3.50 | - | - | - |
| FRS Rasch Score | -0.74 ± 1.55 | - | - | - |
| (Mild, Moderate, Severe, Very Severe) ¶ | (1, 11, 15, 2) | - | - | - |
| Diagnostic certainty (Possible, Probable, Definite) | 3, 22, 6 | - | - | - |
| Gene variants (*C9ORF72*, *GRN*, *MAPT*) | 7, 2, 1 | - | - | - |

Data are presented as mean ± SD. Significant associations are in bold font.
\*Chi-squared test
<sup>†</sup>Independent sample t tests
FRS = Frontotemporal dementia rating scale, ACE-R = Addenbrooke's Cognitive Examination
Revised.
Missing data for <sup>‡</sup>one, <sup>§</sup>three and <sup>¶</sup>two bvFTD cases.

All participants or their responsible caregiver provided written informed consent in accordance with the Declaration of Helsinki. The South Eastern Sydney Local Health District, University of New South Wales (HC12573), and University of Sydney (HREC 10/126 and HE000408) ethics committees approved the study.

### 2.2 Lipid Quantification

Plasma was prepared from whole blood collected into heparin-coated tubes, aliquoted, and stored at -80 □. Lipids were extracted from 50 μL plasma using the methyl-tert-butyl ether/methanol/water protocol,^11^ with 400 pmoles of GluCer(d18:1/12:0) internal standard 5 (#860543, Avanti Polar Lipids) added prior to extraction. Lipids were reconstituted in 100 μL methanol, stored at -30°C, and diluted 1:50 in HPLC mobile phase immediately prior to measurement of GluCer, GalCer, and HexCer by LC-MS/MS.^12^ Full details for lipid extraction and analysis are provided in the Supplementary File.

### 2.3 Statistical Analyses on Lipidomic Data

Lipid levels were ln-transformed to improve normality, after which all lipids except C22:1 GluCer were normally distributed in the Anderson-Darling and/or D’Agostino-Pearson tests. Lipid levels were compared between control and bvFTD groups using two-tailed t-tests, and multiple linear regression adjusting for age and sex. Pearson’s analysis was used to test correlations between lipids and age. Spearman analysis (two-tailed) was used to test correlations between lipids and ACE-R scores or disease duration (which were not normally distributed). The two-stage step-up method of Benjamini, Krieger and Yekutieli was used to correct *P* values for false discovery rate, and corrected *P* values are reported as *Q* values. *Q*<0.05 was considered statistically significant.

### 2.4 MRI acquisition

MRI was acquired on a Philips Achieva 3.0T scanner using a standard 8-channel head coil. Single-shell diffusion weighted imaging (DWI) data were obtained using the following sequences and parameters: two sets with 32 isotropic diffusion directions at a b-value of 1000 s/mm. T1-weighted images were also acquired with the following parameters: repetition time/echo time 2.6/5.8 ms, 200 slices, voxel resolution 1 mm isotropic, in-plane matrix: 256 × 256, flip angle α=8.

### 2.5 MRI preprocessing and analysis

DWI images were pre-processed using standardised MRtrix3^16^ and FSL^17^ pipelines. Preprocessing steps included denoising, and corrections for standard distortion, eddy currents, motion, and field inhomogeneities. Resulting images were visually inspected before normalisation and registration. Fibre density and cross-section (FDC), a measure of white matter fibre bundle integrity, was computed for all participants. Whole-brain imaging analyses using fixel-wise general linear models were conducted to compare (i) FDC changes between bvFTD patients and controls, and (ii) associations between lipid levels and FDC in both bvFTD patients and controls. The statistical threshold for all MRI analyses was initially set at *P*<0.05 family-wise error correction for multiple comparisons, and then repeated at *P*<0.005 uncorrected with a conservative cluster extent threshold of 100 contiguous fixels to minimise Type I error while balancing the risk of Type II errors^18^. Full method details are provided in the Supplementary File.

## 3 Results

### 3.1 Participants

Demographic and clinical characteristics of the participants are shown in Table 1. Age and years of education were higher in the control group, whereas sex distribution did not differ significantly between the groups. ACE-R scores were significantly lower in the bvFTD group. MRI within one year of plasma sampling was available for 20 control and 25 bvFTD cases (Supplementary Table 1).

### 3.2 Plasma glycosphingolipids are lower in bvFTD

Levels of six GalCer and six equivalent GluCer species were quantified in plasma samples (Fig. 1A). Only glycosphingolipids with a typical D-erythro-sphingosine base were quantified, and shorthand notation indicating the length and number of double bonds in the N-acyl chain has been used throughout this manuscript, e.g. C24:1 GluCer refers to GluCer(d18:1/24:1), denoting GluCer with a D-erythro-sphingosine (d18:1) backbone and 24-carbon, monounsaturated N-acyl chain.

**Figure 1.**
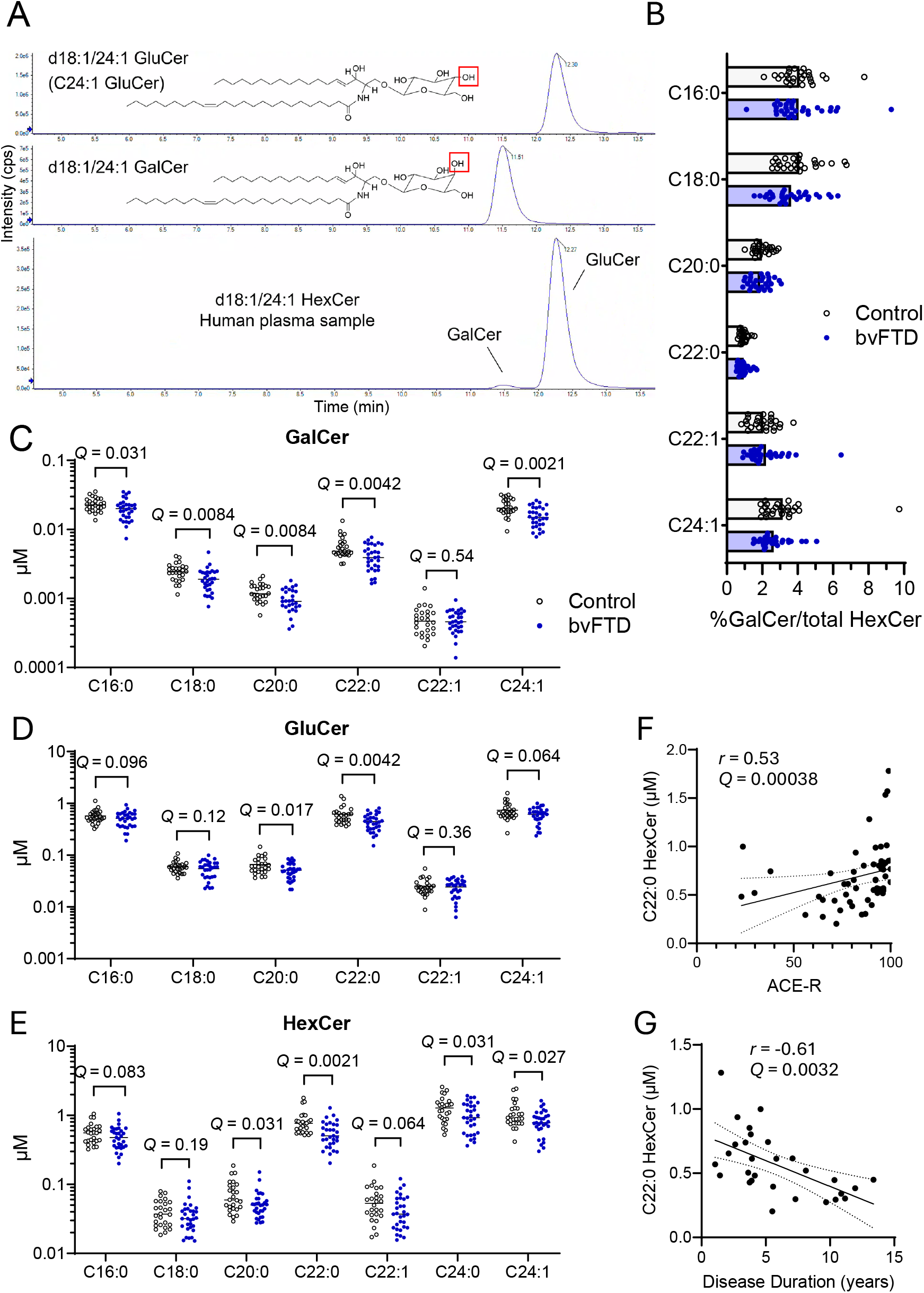
Plasma glycosphingolipid levels in bvFTD and control cases. **(A)** Representative chromatograms showing elution times for C24:1 GluCer (top panel) and C24:1 GalCer (middle panel) standards, and separation of these HexCer isomers in a human plasma sample (lower panel). Structures are shown on the left with red box illustrating the hydroxyl group that differs in chirality between GluCer and GalCer. **(B)** Abundance of GalCer as a percentage of total HexCer for each N-acyl chain length in control and bvFTD plasma samples. Bar shows the mean. **(C)** GalCer, **(D)** GluCer, and **(E)** HexCer concentrations in control and bvFTD plasma samples. **(F, G)** Scatter plots show correlations between plasma levels of C22:0 HexCer and **(F)** ACE-R scores or **(G)** disease duration. Line of best fit and 95% confidence intervals are shown, *r* and *Q* values are derived from Spearman correlation analysis.

In control plasma, HexCer was comprised 96-99% of GluCer, depending on the specific species (Fig. 1B). There was no significant difference in the relative proportion of GalCer to GluCer between bvFTD and controls. After correcting for false discovery rate, levels of five of the six GalCer (C16:0, C18:0, C20:0, C22:0, and C24:1) and two GluCer (C20:0 and C22:0) species were significantly lower in bvFTD compared to control samples (Fig. 1C, D). Since quantification of HexCer with reverse phase LC-MS/MS is more common and simpler than separation of GluCer and GalCer with HILIC-MS/MS, we determined if the changes to GluCer and GalCer were reflected in changes to circulating HexCer. Of seven HexCer species measured, C20:0, C22:0, C24:0, and C24:1 were significantly reduced in bvFTD compared to control plasma (Fig. 1E).

### 3.3 Plasma glycosphingolipids are correlated with cognitive scores and disease duration

Twelve of the 19 lipids measured were positively correlated with ACE-R at *Q*<0.05, and seven were inversely correlated with disease duration (Table 2). C22:0 HexCer showed the strongest correlations with both ACE-R and disease duration (Fig. 1F, G), noting that the correlation to ACE-R scores appeared not to hold for ACE-R scores <50.

**Table 2.**
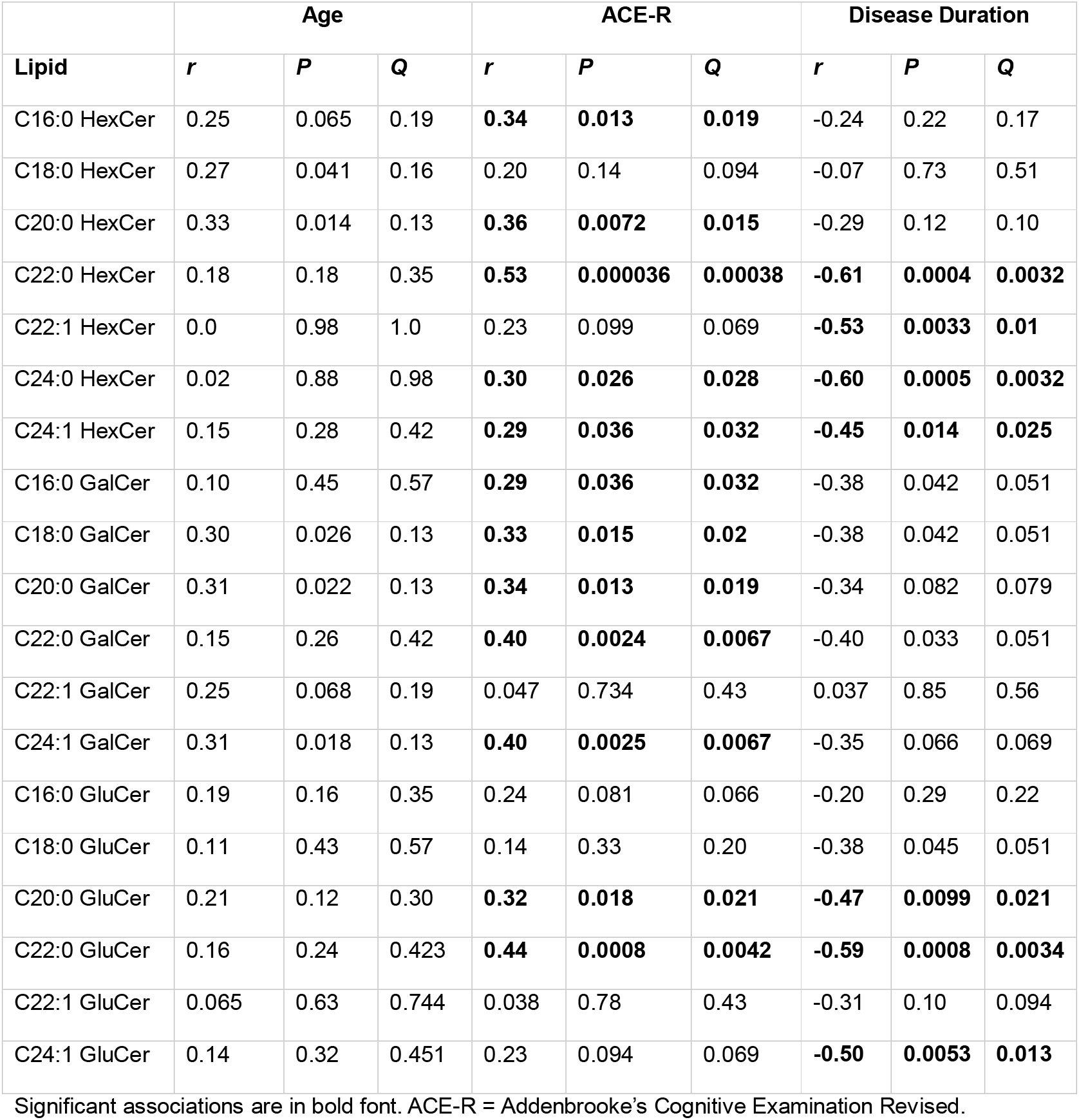
Univariate associations of lipids with age, ACE-R scores and disease duration.

| Lipid | Age |  |  | ACE-R |  |  | Disease Duration |  |  |
| --- | --- | --- | --- | --- | --- | --- | --- | --- | --- |
|  | *r* | *P* | *Q* | *r* | *P* | *Q* | *r* | *P* | *Q* |
| C16:0 HexCer | 0.25 | 0.065 | 0.19 | **0.34** | **0.013** | **0.019** | -0.24 | 0.22 | 0.17 |
| C18:0 HexCer | 0.27 | 0.041 | 0.16 | 0.20 | 0.14 | 0.094 | -0.07 | 0.73 | 0.51 |
| C20:0 HexCer | 0.33 | 0.014 | 0.13 | **0.36** | **0.0072** | **0.015** | -0.29 | 0.12 | 0.10 |
| C22:0 HexCer | 0.18 | 0.18 | 0.35 | **0.53** | **0.000036** | **0.00038** | **-0.61** | **0.0004** | **0.0032** |
| C22:1 HexCer | 0.0 | 0.98 | 1.0 | 0.23 | 0.099 | 0.069 | **-0.53** | **0.0033** | **0.01** |
| C24:0 HexCer | 0.02 | 0.88 | 0.98 | **0.30** | **0.026** | **0.028** | **-0.60** | **0.0005** | **0.0032** |
| C24:1 HexCer | 0.15 | 0.28 | 0.42 | **0.29** | **0.036** | **0.032** | **-0.45** | **0.014** | **0.025** |
| C16:0 GalCer | 0.10 | 0.45 | 0.57 | **0.29** | **0.036** | **0.032** | -0.38 | 0.042 | 0.051 |
| C18:0 GalCer | 0.30 | 0.026 | 0.13 | **0.33** | **0.015** | **0.02** | -0.38 | 0.042 | 0.051 |
| C20:0 GalCer | 0.31 | 0.022 | 0.13 | **0.34** | **0.013** | **0.019** | -0.34 | 0.082 | 0.079 |
| C22:0 GalCer | 0.15 | 0.26 | 0.42 | **0.40** | **0.0024** | **0.0067** | -0.40 | 0.033 | 0.051 |
| C22:1 GalCer | 0.25 | 0.068 | 0.19 | 0.047 | 0.734 | 0.43 | 0.037 | 0.85 | 0.56 |
| C24:1 GalCer | 0.31 | 0.018 | 0.13 | **0.40** | **0.0025** | **0.0067** | -0.35 | 0.066 | 0.069 |
| C16:0 GluCer | 0.19 | 0.16 | 0.35 | 0.24 | 0.081 | 0.066 | -0.20 | 0.29 | 0.22 |
| C18:0 GluCer | 0.11 | 0.43 | 0.57 | 0.14 | 0.33 | 0.20 | -0.38 | 0.045 | 0.051 |
| C20:0 GluCer | 0.21 | 0.12 | 0.30 | **0.32** | **0.018** | **0.021** | **-0.47** | **0.0099** | **0.021** |
| C22:0 GluCer | 0.16 | 0.24 | 0.423 | **0.44** | **0.0008** | **0.0042** | **-0.59** | **0.0008** | **0.0034** |
| C22:1 GluCer | 0.065 | 0.63 | 0.744 | 0.038 | 0.78 | 0.43 | -0.31 | 0.10 | 0.094 |
| C24:1 GluCer | 0.14 | 0.32 | 0.451 | 0.23 | 0.094 | 0.069 | **-0.50** | **0.0053** | **0.013** |

Significant associations are in bold font. ACE-R = Addenbrooke's Cognitive Examination Revised.

No lipids were significantly associated with age or sex at *Q*<0.05, however at *P*<0.05, five lipids were positively correlated with age (Table 2) and three were higher in females compared to males (Supplementary Table 2). We therefore tested associations between lipids and bvFTD after adjusting for age and sex, finding that C22:0 GalCer (*F*(1,52)=7.4, *P*=0.009), C24:1 GalCer (*F*(1,52)=8.8, *P*=0.005), C22:0 GluCer (*F*(1,52)=7.9, *P*=0.007), and C22:0 HexCer (*F*(1,52)=9.7, *P*=0.003) were significantly lower in bvFTD compared to control subjects (all *Q*=0.035) (Supplementary Table 3).

### 3.4 Plasma HexCer is correlated with white matter tract integrity in bvFTD

Compared to controls, bvFTD patients exhibited significantly reduced FDC in the frontal commissural fibres of the corpus callosum and forceps minor, extending posteriorly along the superior longitudinal fasciculi bilaterally. Reduced FDC was also observed in the frontotemporal association fibres of the uncinate and inferior longitudinal fasciculi, as well as in the cerebellopontine fibres and projections to the superior frontal gyrus in the left hemisphere (Fig. 2A).

**Figure 2.**
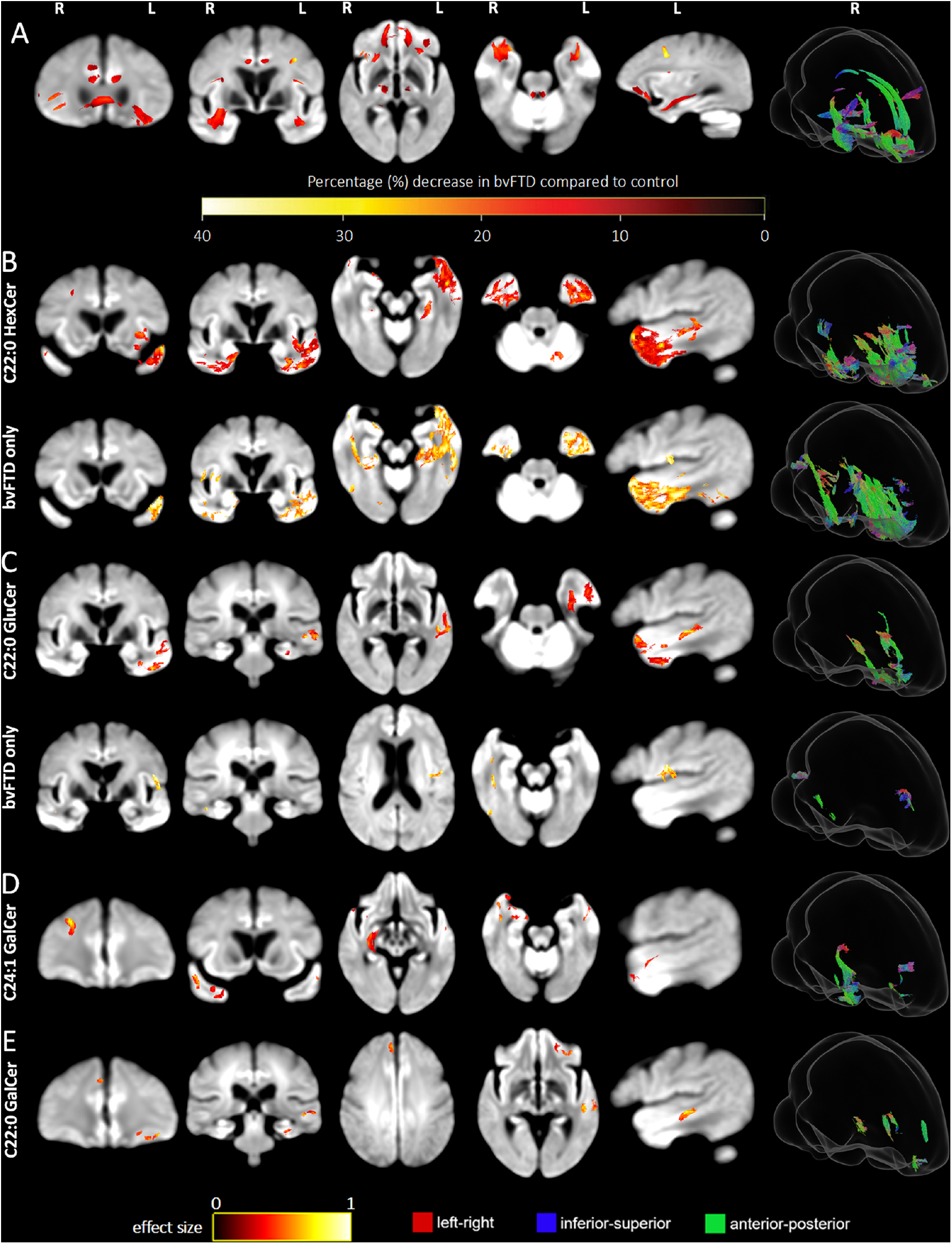
Plasma lipid levels are correlated with FDC reductions in frontotemporal white matter. Brain maps show white matter tracts for which **(A)** FDC is significantly reduced in bvFTD (*n* = 25) compared to control (*n* = 20) cases at *P*<0.05 corrected for family-wise error. **(B-E)** FDC is correlated with levels of **(B)** C22:0 HexCer, **(C)** C22:0 GluCer, **(D)** C24:1 GalCer, and **(E)** C22:0 GalCer at *P*<0.005, uncorrected for family-wise error. The correlations for the bvFTD group alone are also shown for C22:0 HexCer **(B)** and C22:0 GluCer **(C)**. Colours indicate the percentage decrease compared to controls in **(A)** or effect size **(B-E)**. In glass brain images on the right, colour indicates tract direction, red: left-right, green: anterior-posterior, blue: superior-inferior. R: right, L: left.

Next, we analysed the associations between lipids and white matter for the four lipids that demonstrated significant differences between bvFTD patients and controls, after adjusting for age and sex. All four lipids showed positive correlations with the FDC of frontotemporal white matter fibres in the full cohort (*P*<0.005 uncorrected for family-wise error, Fig. 2B-E and Supplementary Figure 1). C22:0 HexCer was associated with reduced FDC in the inferior longitudinal fasciculus fibres in the left inferior, middle and superior temporal left lobe. Further changes were seen in frontal association fibres, middle commissural fibres, right inferior longitudinal fasciculus, left insula, left uncinate, and left cerebellar peduncles (Fig. 2B). Considering only the bvFTD cases, the effect size of the association was stronger and observed in similar regions as seen with the full cohort. C22:0 GluCer was associated with FDC reductions in similar fibre bundles, albeit only in the left temporal lobe and to a lesser extent (Fig. 2C). Considering only the bvFTD cases, the associations were minimal, involving the left middle temporal tracts. There were no significant associations with C22:0 HexCer or C22:0 GluCer when analysing only the control group.

The associations between C24:1 or C22:0 GalCer and FDC were more limited. C24:1 GalCer was associated with mostly right lateralised FDC reductions in the inferior longitudinal fasciculus and superior frontal and middle commissural fibres (Fig. 2D). C22:0 GalCer associated with more sparse FDC reductions in the left inferior longitudinal fasciculus and frontal white matter fibres in both hemispheres (Fig. 2E). These correlations were absent when analysing the bvFTD and control groups separately.

## 4 Discussion

This study establishes that levels of very long chain (C20-C24) HexCer species are lower in plasma of bvFTD compared to control subjects, positively correlated with cognitive performance, and inversely correlated with disease duration. In particular, both GluCer and GalCer with a C22:0 N-acyl chain were significantly reduced in bvFTD after adjusting for age and sex, and this was confirmed by measuring their composite HexCer using a different LC-MS/MS method and instrument. Plasma C22:0 HexCer and GluCer levels were associated with reductions in density and cross-section of frontal and temporal association fibres in the white matter, urging further investigation into the utility of these lipids as blood biomarkers of white matter degeneration and disease severity in bvFTD.

Plasma levels of both GluCer and GalCer were lower in bvFTD, and C22:0 GluCer was more strongly correlated with FDC of temporal and frontal white matter tracts than C22:0 GalCer. This implies that reduced glycosphingolipid levels in bvFTD are not directly attributable to demyelination, despite the abundance of GalCer in myelin,^9,12^ and may instead reflect changes to peripheral lipid metabolism. The observation that HexCer levels correlate more strongly with white matter integrity than either GluCer or GalCer may result from more robust quantification of HexCer with reverse phase compared to HILIC LC-MS/MS. Furthermore, since C22:0 HexCer and C22:0 GluCer were correlated with white matter integrity in bvFTD but not control cases, these associations cannot be attributed to the differences in lipid levels between the control and bvFTD groups.

There have been few lipidomic analyses of FTD blood samples. A prior study reported increased levels of triglyceride and lysophosphatidylcholine species, and decreased acylcarnitines and cardiolipins, in bvFTD serum samples.^19^ In plasma, total triglycerides were increased, while phosphatidylserine and phosphatidylglycerol were decreased.^20^ Reductions in HexCer were not noted, likely a consequence of using less quantitative untargeted methods and reporting lipid class totals, which do not necessarily reflect the abundance of individual lipid species. Plasma HexCer levels were either unaltered or higher than controls in two large Alzheimer’s Disease cohorts,^21^ while levels of multiple HexCer species were higher in plasma and serum of people with progressive multiple sclerosis,^22,23^ in which serum C22:0 HexCer and plasma C20:0 HexCer were positively correlated with retinal nerve fibre atrophy^22^ and brain atrophy,^23^ respectively. This suggests that reduced plasma HexCer levels in bvFTD are somewhat specific to the disease and not directly associated with demyelination or neurodegeneration.

Hexosylceramides with C20-C24 N-acyl chains were more significantly decreased than those with C16 or C18 chains in bvFTD plasma. Circulating lipids are derived in large part from the liver, and C20-C24 sphingolipids are produced by ceramide synthase 2 (CerS2), which is highly expressed in the liver.^24^ Hepatocyte CerS2 is essential for insulin sensitivity and metabolic health,^25,26^ while CerS2 in oligodendrocytes is essential for myelin stability.^12^ In addition to myelin deficits, bvFTD is associated with increased BMI and plasma triglycerides, insulin resistance, and higher incidence of diabetes.^27^ Reduced levels of C20-C24 glycosphingolipids might therefore reflect the metabolic phenotype of bvFTD. However, a selective reduction of CerS2 activity in bvFTD seems unlikely, given that mean levels of C16:0 and C18:0 glycosphingolipids, which are synthesized by other ceramide synthases,^24^ were also lower (statistically significant for GalCer). The possibility of altered peripheral sphingolipid homeostasis in bvFTD should be investigated in future studies by quantifying the metabolically-related sphingolipids ceramide, sphingomyelin, and lactosylceramide.

Another possibility is that bvFTD is characterised by reduced availability of very long chain fatty acids, which are substrates for sphingolipid synthesis.^24^

Given the higher average BMI^27^ and triglycerides^20^ in people with bvFTD, a limitation of our study was the absence of BMI data. Hexosylceramides show modest inverse correlations with BMI in large cohorts,^21,28^ however reduced HexCer levels in bvFTD are unlikely to be solely attributed to BMI, due to their associations with disease duration, cognitive scores, and white matter integrity.

Current dementia staging models suggest that white matter changes occur before grey matter atrophy and clinical symptoms^29^, while longitudinal imaging studies have shown that brain atrophy in bvFTD spreads posteriorly and towards the right hemisphere as the disease progresses.^30^ Herein, we found that plasma C22:0 HexCer and GalCer were strongly associated with white matter loss in typical baseline brain atrophy regions in bvFTD. Since these changes were also linked to disease duration and severity, quantification of plasma lipids could serve as a non-invasive, cheap, and easy-to-deploy peripheral biomarker for early white matter degeneration with clinical significance.

In conclusion, lower levels of C22:0 HexCer and related glycosphingolipids are indicative of white matter degeneration, longer disease duration and cognitive impairment in bvFTD. Our findings should be validated in a distinct cohort, determining if levels of these lipids are reduced in other forms of FTD or related neurological diseases. Longitudinal studies with larger sample sizes are needed to test the accuracy and predictive value of these lipid markers for staging bvFTD *in-vivo*, which would aid the recruitment and monitoring of patients in drug trials for disease-modifying therapies.

## Acknowledgements

We thank all the participants and their carers for their time and contribution to this study. We gratefully acknowledge the infrastructure and subsidised access provided by the Sydney Mass Spectrometry, Sydney Imaging, and the Sydney Informatics Hub core facilities at the University of Sydney.

## Data Availability

The data supporting the conclusions of this manuscript are available upon reasonable request to the corresponding author.

