## Supplementary File for "Reduced plasma hexosylceramides in frontotemporal dementia are a biomarker of white matter integrity"

**Supplementary Methods**

**Lipid Extraction**

Lipids were extracted from 50 µL of plasma using the methyl tert-butyl ether (MTBE)/methanol/water protocol as described.^1^ Plasma was combined with 850 µL MTBE and 250 µL methanol containing 400 pmoles GluCer(d18:1/12:0) internal standard (#860543, Avanti Polar Lipids). After sonicating at 4℃ for 30 min, 162 µL of water was added, samples were vortexed, then centrifuged at 2000 × *g* for 5 min. The upper organic phase was collected in 5 mL glass tubes. The lower phase was extracted twice more with 500 µL MTBE and 150 µL methanol, followed by sonication and phase separation with 125 µL water. The organic phases were combined and dried in a Savant SC210 SpeedVac (ThermoFisher Scientific). Lipids were reconstituted in 100 µL methanol and stored at -30°C.

**Separation and Quantification of GluCer and GalCer**

GluCer and GalCer were analysed by multiple reaction monitoring on a Sciex 6500+ QTRAP mass spectrometer coupled to a Shimadzu Nexera HPLC as previously described.^2^ Lipids were resolved on a 2.1 x 150 mm Agilent InfinityLab Poroshell 120 HILIC-Z HPLC column (2.7 µm pore size), with column oven at 30°C and an isocratic mobile phase comprising 97% acetonitrile, 2% methanol, 0.75% water, 0.25% formic acid, and 2.5 mM ammonium formate. Lipid extracts in methanol were diluted 1:50 into the HPLC mobile phase without methanol immediately prior to analysis. Run time was 20 min, and flow rate 0.1 mL/min. Precursor *m/z* was the relevant [M+H]^+^ ion, and product *m/z* was 264.3. Source parameters were spray voltage: 4000V, temperature: 225°C, ion source gas 1 and 2: 20 psi, curtain gas: 40 psi and CAD gas: 12 (arbitrary units). The entrance potential (EP) was 10V, the collision energy (CE) 45V and collision cell exit potential (CXP) 12V. Sample order was randomised, and SciexOS (version 1.7) was used for peak integration. Separation of GalCer and GluCer was confirmed using GalCer(d18:1/24:1) and GluCer(d18:1/24:1) (C24:1 GalCer and C24:1 GluCer) standards (Avanti Polar Lipids, #860432 and #860549). The amount of each lipid was calculated as the ratio to the internal standard, multiplied by 400 pmoles.

**Quantification of HexCer**

HexCer species were detected using multiple reaction monitoring on a TSQ Altis triple quadrupole mass spectrometer coupled to a Vanquish HPLC (Thermo Scientific). Lipids were resolved on a 3x150 mm Agilent ZORBAX Eclipse XDB-C8 column (5 µm pore size), with column oven at 30°C using an isocratic solvent comprising 2 mM ammonium formate and 0.2% formic acid in 100% methanol. Run time was 10 min at a flow rate of 0.4 mL/min. Precursor *m/z* was the relevant [M+H]^+^ ion, and product *m/z* was 264.3. Source parameters were spray voltage: 4500V, sheath gas: 8.5, aux gas: 10, sweep gas: 10, ion transfer tube temperature: 275°C, vaporizer temperature: 150°C. The CE was 34V and radio frequency (RF) lens was 43V. TraceFinder (version 5.1) was used for peak integration.

**MRI Data Analysis**

MRI preprocessing included: denoising,^3,4^ Synb0 distortion correction using the T1,^5^ eddy-current and motion correction^6,7^ and bias field correction.^5^ The quality of each pre-processed DWI was visually inspected before proceeding to group pre-processing. Global intensity normalization was then carried out across all subjects.^8^ A group-average white matter response was then computed for estimating fiber orientation distributions (FODs).^9^ DWI data was re-sampled to an isotropic voxel size of 1.25 mm,^10^ and FODs were computed with Single-Shell, 3-Tissue Constrained Spherical Deconvolution.^11^ To improve registration errors and detection sensitivity in downstream analyses, spatial normalisation was achieved by generating a study-specific unbiased FOD population template with an iterative registration and averaging approach.^12^ Each patient’s relevant FOD image was subsequently registered to the template.

Using the MRtrix3^13,14^ analysis package, three metrics of fibre bundle integrity were obtained for each subject across all white matter fixels (individual fibre populations within a voxel^15^). Fibre density (FD) was calculated as the volume of the intra-axonal compartment within fixels.^8^ Fibre cross-section (FC) was calculated as difference in fibre-bundle diameter between the subject and template. FDC was calculated as the FD multiplied by the FC. Twenty million streamlines were generated then subsequently filtered to two million using the spherical-deconvolution informed filtering of tractograms algorithm to reduce reconstruction biases.^16^

**Statistical Analyses on MRI Data**

White matter integrity was analysed with fixel-wise general linear models comparing the FDC metric between patients and controls. Connectivity-based fixel enhancement was applied to smooth the fixel metrics and assist with statistical inference of the template tractogram.^17^ Multiple comparisons correction was conducted using non-parametric permutation to assign family-wise error corrected p-values.^18,19^ Separate fixel-wise general linear models were used to test associations between each lipid of interest and FDC in all (control and bvFTD) subjects together. All subjects were included in the models to increase the number and spread of datapoints and thus increase the power to detect associations. Standardised effect sizes were calculated with the Cohen’s d.^20^ All fixel-wise statistical analyses were carried out using MRtrix3 software.^13,14^

**Supplementary Results**

**Supplementary Table 1** **Demographic and clinical index data in bvFTD and control cases with matching MRI.**

| **Characteristic** | **bvFTD (n=25)** | **Control (n=20)** | **Statistic** | ***P*** |
| --- | --- | --- | --- | --- |
| Sex (Female:Male) | 12:13 | 9:11 | 0.04^*^ | 0.84 |
| Age (years) | 63.61 ± 6.40 | 69.64 ± 6.70 | 3.06^†^ | **0.004** |
| Education (years) | 12.16 ± 3.09 | 14.29 ± 2.41 | 2.59 ^†^ | **0.013** |
| ACE-R | 72.66 ± 19.35 | 95.91 ± 2.60 | 5.94^†^ | **<0.001** |
| Disease duration (years) | 5.13 ± 3.10 | - | - | - |
| FRS Rasch Score  (Mild:Moderate:Severe:Very severe) | -0.51 ± 1.46 (1:11:13:0) | - | - | - |
| Diagnostic certainty (Possible:Probable:Definite) | 3:18:4 | - | - | - |
| Gene variants (C9ORF72:GRN:MAPT) | 4:2:1 |  |  |  |

Data are presented as mean ± SD. Significant differences are in bold font.

^*^ Chi-squared test
^†^ Independent sample t tests;
FRS = Frontotemporal dementia rating scale, ACE-R = Addenbrooke’s Cognitive Examination Revised.

**Supplementary Table 2 Univariate associations of lipids with sex.**

| **Lipid** | **Male** | **Female** | ***t*** | ***P*** | ***Q*** |
| --- | --- | --- | --- | --- | --- |
| C16:0 HexCer | 0.53 ± 0.18 | 0.53 ± 0.21 | 0.35 | 0.73 | 0.92 |
| C18:0 HexCer | 0.038 ± 0.017 | 0.04 ± 0.022 | 0.072 | 0.94 | 0.99 |
| C20:0 HexCer | 0.059 ± 0.027 | 0.067 ± 0.041 | 0.55 | 0.58 | 0.83 |
| C22:0 HexCer | 0.62 ± 0.28 | 0.75 ± 0.34 | 1.76 | 0.083 | 0.33 |
| C22:1 HexCer | **0.042 ± 0.025** | **0.062 ± 0.035** | **2.66** | **0.01** | **0.1** |
| C24:0 HexCer | **0.98 ± 0.47** | **1.33 ± 0.51** | **2.88** | **0.0056** | **0.1** |
| C24:1 HexCer | 0.87 ± 0.33 | 1.03 ± 0.5 | 1.19 | 0.24 | 0.48 |
| C16:0 GalCer | 0.021 ± 0.0068 | 0.022 ± 0.0058 | 0.61 | 0.55 | 0.83 |
| C18:0 GalCer | 0.0021 ± 0.00073 | 0.0023 ± 0.00084 | 1.13 | 0.26 | 0.48 |
| C20:0 GalCer | **0.001 ± 0.00031** | **0.0012 ± 0.00043** | **2.06** | **0.045** | **0.29** |
| C22:0 GalCer | 0.0045 ± 0.0019 | 0.0053 ± 0.0023 | 1.59 | 0.12 | 0.34 |
| C22:1 GalCer | 0.00049 ± 0.00016 | 0.00051 ± 0.00025 | 0.18 | 0.86 | 0.95 |
| C24:1 GalCer | 0.017 ± 0.0052 | 0.02 ± 0.0069 | 1.94 | 0.06 | 0.29 |
| C16:0 GluCer | 0.54 ± 0.16 | 0.54 ± 0.19 | 0.2 | 0.84 | 0.95 |
| C18:0 GluCer | 0.054 ± 0.019 | 0.06 ± 0.017 | 1.4 | 0.17 | 0.41 |
| C20:0 GluCer | 0.053 ± 0.019 | 0.063 ± 0.025 | 1.64 | 0.11 | 0.32 |
| C22:0 GluCer | 0.49 ± 0.2 | 0.56 ± 0.24 | 1.29 | 0.2 | 0.45 |
| C22:1 GluCer | 0.024 ± 0.008 | 0.025 ± 0.01 | 0.34 | 0.74 | 0.92 |
| C24:1 GluCer | 0.63 ± 0.21 | 0.71 ± 0.27 | 1.01 | 0.32 | 0.53 |

Data are presented as mean ± SD concentration (µM). Significant associations for independent t-test at *P*<0.05 are in bold font.

**Supplementary Table 3 Association of lipids with bvFTD in linear regression adjusted for age and sex.**

|  | **bvFTD** | | | **Sex** | | **Age** | |
| --- | --- | --- | --- | --- | --- | --- | --- |
| **Lipid** | **F** | ***P*** | ***Q*** | **F** | ***P*** | **F** | ***P*** |
| C16:0 GalCer | 3.35 | 0.073 | 0.12 | 0.049 | 0.83 | 0.013 | 0.91 |
| C18:0 GalCer | **4.2** | **0.046** | 0.12 | 0.9 | 0.35 | 2.5 | 0.12 |
| C20:0 GalCer | **4.44** | **0.04** | 0.12 | **4.15** | **0.047** | 2.86 | 0.097 |
| C22:0 GalCer | **7.41** | **0.0088** | **0.035** | 1.23 | 0.26 | 0.097 | 0.76 |
| C22:1 GalCer | 0.3 | 0.59 | 0.55 | 0.15 | 0.7 | 3.72 | 0.059 |
| C24:1 GalCer | **8.78** | **0.0046** | **0.035** | 3.02 | 0.088 | 2.65 | 0.11 |
| C16:0 HexCer | 1.19 | 0.28 | 0.32 | 0.19 | 0.66 | 1.83 | 0.18 |
| C18:0 HexCer | 0.16 | 0.69 | 0.6 | 0.022 | 0.88 | 3.25 | 0.077 |
| C20:0 HexCer | 2.01 | 0.16 | 0.21 | 0.23 | 0.63 | 3.8 | 0.057 |
| C22:0 HexCer | **9.65** | **0.0031** | **0.035** | 2.02 | 0.16 | 0.21 | 0.65 |
| C22:1 HexCer | 2.47 | 0.12 | 0.17 | **5.36** | **0.025** | 0.12 | 0.73 |
| C24:0 HexCer | 3.63 | 0.062 | 0.12 | **6.22** | **0.016** | 0.1 | 0.75 |
| C24:1 HexCer | 3.31 | 0.075 | 0.12 | 0.66 | 0.42 | 0.25 | 0.62 |
| C16:0 GluCer | 1.1 | 0.3 | 0.32 | 0.15 | 0.7 | 0.89 | 0.35 |
| C18:0 GluCer | 0.905 | 0.35 | 0.35 | 1.51 | 0.23 | 0.26 | 0.61 |
| C20:0 GluCer | 3.53 | 0.066 | 0.12 | 1.94 | 0.17 | 1.0 | 0.32 |
| C22:0 GluCer | **7.9** | **0.0069** | **0.035** | 0.84 | 0.36 | 0.087 | 0.77 |
| C22:1 GluCer | 0.068 | 0.8 | 0.66 | 0.035 | 0.85 | 0.13 | 0.72 |
| C24:1 GluCer | 1.51 | 0.22 | 0.27 | 0.43 | 0.52 | 0.34 | 0.56 |

Significant associations for *P*<0.05 are in bold font. No lipids were significantly affected by age or sex at *Q*<0.05, therefore *Q* values are not shown for these variables.

**Supplementary Figure 1 (animation)**

Rotating glass brain images show white matter tracts for which FDC is correlated with levels of (A) C22:0 HexCer, (B) C22:0 GluCer, (C) C24:1 GalCer, and (D) C22:0 GalCer at p<0.005, uncorrected for family-wise error. Dashes indicate no significant findings at that level of significance. Colour indicates tract direction, red: left-right, green: anterior-posterior, blue: superior-inferior. The left column shows results for all subjects while the right column shows results for the bvFTD group only.
